# Co-option of mitochondrial nucleic acid sensing pathways by HSV-1 UL12.5 for reactivation from latent Infection

**DOI:** 10.1101/2024.07.06.601241

**Authors:** Sean R. Cuddy, Matthew E. Flores, Patryk A. Krakowiak, Abigail L. Whitford, Sara A. Dochnal, Aleksandra Babnis, Tsuyoshi Miyake, Marco Tigano, Daniel A. Engel, Anna. R Cliffe

**Affiliations:** Neuroscience Graduate Program, University of Virginia, Charlottesville, VA, 22908; Department of Microbiology, Immunology and Cancer Biology, University of Virginia, Charlottesville, VA, 22908; Department of Pathology and Genomic Medicine, Thomas Jefferson University, 1020 Locust Street, Philadelphia 19107

**Author notes:** These authors contributed equally (Sean R. Cuddy, Matthew E. Flores, Patryk A. Krakowiak).

## Abstract

Although viruses subvert innate immune pathways for their replication, there is evidence they can also co-opt anti-viral responses for their benefit. The ubiquitous human pathogen, Herpes Simplex Virus-1 (HSV-1), encodes a protein (UL12.5) that induces the release of mitochondrial nucleic acid into the cytosol, which activates immune sensing pathways and reduces productive replication in non-neuronal cells. HSV-1 establishes latency in neurons and can reactivate to cause disease. We found that UL12.5 is required for HSV-1 reactivation in neurons and acts to directly promote viral lytic gene expression during initial exit from latency. Further, the direct activation of innate immune sensing pathways triggered HSV reactivation and compensated for a lack of UL12.5. Finally, we found that the induction of HSV-1 lytic genes during reactivation required intact RNA and DNA sensing pathways, demonstrating that HSV-1 can both respond to and active antiviral nucleic acid sensing pathways to reactivate from a latent infection.

## Introduction

Viruses and their hosts continually antagonize each other in a battle between host immunity and viral persistence. The success of persistent viruses can be attributed to the subversion or co-option of host innate immune pathways mediated by pattern-recognition receptors (PRRs) that sense PAMPs (Pathogen-Associated Molecular Patterns)^1^. Although PPRs recognize pathogen-derived molecules, they can also recognize misplaced host molecular patterns, including cytosolic mitochondrial DNA (mtDNA), to result in the activation of innate immune responses ^2^. Somewhat paradoxically, flaviviruses, influenza A virus, herpesviruses, and others express proteins that specifically induce mtDNA stress that is associated with mtDNA release and subsequent activation of innate immune pathways ^3–6^. However, it is unknown whether there is a beneficial role mtDNA release in the context of viral infection.

Of the PRRs that respond to viral infection and mtDNA, cGAS (cyclic GMP-AMP synthase) and STING (stimulator of interferon genes) mediate a cytosolic DNA-sensing pathway that is one of the most evolutionarily conserved nucleic acid-sensing pathways found in mammals and plays a central role in the cellular response to infection ^7–9^. Upon sensing cytosolic DNA, cGAS dimerizes and produces the secondary messenger 2’-3’-cyclic GMP-AMP (2’-3’-cGAMP) that induces STING’s activation and subsequent translocation from the endoplasmic reticulum to the Golgi. mtRNA released into the cytosol can also be sensed by the RNA-ligands RIG-I (retinoic acid-inducible gene I) or MDA5 (melanoma differentiation-associated protein 5), resulting in the activation of MAVS (mitochondrial antiviral-signaling protein) ^10,11^. Depending on the cell type, STING or MAVS activation typically leads to an inflammatory response mediated by nuclear factor κB (NF-κB) and interferon regulatory factor (IRF) family of proteins, and c-Jun N-terminal kinase (JNK) to result in transcription of genes encoding type I interferon (IFN), additional cytokines and chemokines, along with some antiviral genes that are directly induced via this pathway ^12,13^.

As a DNA virus that has co-evolved with humans, it is unsurprising that herpes simplex virus-1 (HSV-1) has concurrently evolved with nucleic acid-sensing pathways. HSV-1 is well known to activate the cGAS-STING pathway upon infection ^9,14–16^ and also encodes many viral proteins that inhibit or modulate the cGAS-STING pathway to counteract the resulting innate immune response ^17–22^. During lytic replication, viral gene expression occurs as an ordered cascade: viral immediate-early (IE) mRNAs dependent on host and viral transcription factors and encode proteins that promote the expression of viral early (E) mRNA, followed by the transcription of viral late (L) genes that are dependent upon viral DNA (vDNA) replication. The breadth and redundancy of viral proteins from each gene class that either disrupt or inhibit the host immune response to cytosolic DNA underscore the importance of modulating these pathways to promote viral viability. It is, therefore, puzzling that HSV encodes a protein, UL12.5, that actively induces an innate immune response ^6,23^.

During lytic infection, the outcome of the expression of UL12.5, a mitochondria-targeted protein, is a restriction of viral replication in cells that can both produce and respond to type I IFN ^6^. As a truncated isoform of the nuclear-resident nuclease UL12, UL12.5 localizes to the mitochondria ^24–27^ and, via mechanisms that are not fully understood, induces the formation of nucleoids that bleb off from mitochondria via the endocytic pathway, followed by rupture, to result in mtDNA release into the cytosol and activation of cGAS ^28^. Ultimately, this process results in a depletion of mtDNA ^27^. UL12.5 expression also results in the depletion of mitochondrial RNA (mtRNA) into the cytosol via an unknown mechanism ^27^. Hence, UL12.5 expression enhances anti-viral immune responses and decreases HSV-1 replication ^6^. Given that UL12.5 is, therefore, not essential for viral replication and even limits replication ^6,29^, it is unclear why HSV-1 has evolved to maintain this function.

The previous studies investigating the impact of UL12.5 on HSV-1 replication were performed in mitotic cells during lytic infection ^6,23^. However, HSV-1 also establishes a latent infection in post-mitotic neurons, from which the virus can reactivate to cause disease. As a highly specialized terminally differentiated cell type, the consequence of UL12.5 expression in neurons is unknown. We hypothesized that UL12.5 plays an important function following HSV-1 infection of neurons. HSV-1 can productively replicate in neurons, and there is evidence that productive replication in neurons occurs as part of the acute phase of the disease to maximize the pool of latently infected cells ^30^. Following the acute infection, HSV-1 establishes a long-term latent infection of neurons. Periodically, HSV re-initiates lytic gene expression in response to certain stimuli, including cell stress and inflammatory neuronal hyperexcitability ^31,32^.

HSV-1 reactivation occurs in a biphasic manner. Phase I consists of a burst of lytic gene expression from all three gene classes (IE, E, L) independent of DNA replication, while the subsequent Phase II recapitulates the ordered viral gene cascade observed in lytic replication and results in the production of infectious virus ^32–37^. Phase I gene expression is thought to be independent of the synthesis of viral proteins, and so far, no viral protein is known to function in Phase I to promote lytic gene expression ^33^. While Phase I may not necessarily progress to Phase II, Phase I represents a critical stage in initiating the reactivation program ^32,33^. Although latency is generally asymptomatic, reactivation can be associated with significant clinical morbidities, including herpes simplex keratitis, encephalitis, and mucosal lesions. The growing evidence for an association between HSV-1 infection and neurodegenerative diseases, however, suggests that latency may also contribute to neurodegenerative diseases potentially characterized by inflammation ^38–47^.

The absence of viral proteins during latency necessitates that HSV exploits existing neuronal signaling pathways to initiate and progress reactivation. Several stimuli have been identified that trigger reactivation, including nerve growth factor (NGF) deprivation ^48,49^, DNA damage ^50^, and neuronal hyperexcitability ^32^ with a notable convergence on the neuronal stress pathway mediated by dual leucine zipper kinase (DLK) and JNK ^32,34,36,37,50,51^. Our recent work has identified interleukin-1β (IL-1β), a major inflammatory cytokine, in triggering HSV reactivation via hyperexcitability ^32^. This finding demonstrated the ability of HSV to co-opt an immune pathway to support reactivation from peripheral neurons. However, whether additional innate immune pathways also support HSV-1 reactivation is unknown. Notably, the downstream consequences of many innate sensing pathways in neurons are not well elucidated. Therefore, we investigated the consequence of UL12.5 expression in neurons and its impact on HSV-1 latency and reactivation. We found that while UL12.5 expression did activate STING, we could not detect increased expression of *Ifn* genes nor an impact on HSV-1 replication. Surprisingly, we did find that UL12.5 was required for HSV-1 reactivation and acted to promote Phase I gene expression, placing UL12.5 as the first viral protein identified to be expressed in Phase I and promote lytic gene expression. Further, reactivation was induced by activating innate sensing pathways and inhibited by the co-depletion of MAVS and STING. Therefore, we have identified a novel mechanism by which HSV-1 utilizes host innate immune signaling pathways to promote lytic gene expression during reactivation.

## Results

### UL12.5 expression does not affect lytic replication in primary neurons

So far, the consequences of UL12.5 expression on viral replication have been investigated exclusively in mitotic cells, such as fibroblasts. However, HSV-1 is a neurotropic virus that can undergo lytic replication in neurons and establishes a latent infection that is capable of reactivating. Therefore, we sought to determine whether UL12.5 was required for lytic replication or latent infection in peripheral neurons. To this end, we isolated primary neurons from the superior cervical ganglia (SCG) of postnatal mice and infected them with a virus lacking UL12.5 expression ^29^. Because UL12.5 is co-linear with the essential *UL12* open-reading frame (ORF), we used the previously described KOS-UL98 virus, in which the entire *UL12* ORF is replaced with *UL98*, an ortholog from cytomegalovirus that maintains the alkaline nuclease function of UL12 but lacks the ability to deplete mitochondrial nucleic acids. Compared to the control virus KOS-SPA, we found no difference in viral replication in primary neurons based on assays for both infectious virus and vDNA replication (Figure 1A & B). This is in contrast to what we observed in primary murine dermal fibroblasts (DFs) isolated from the same mice, where the KOS-UL98 virus replicated to higher levels than the wild-type SPA virus (Figure 1C). Therefore, UL12.5 is neither detrimental nor required for HSV-1 replication in peripheral neurons.

**Figure 1.**
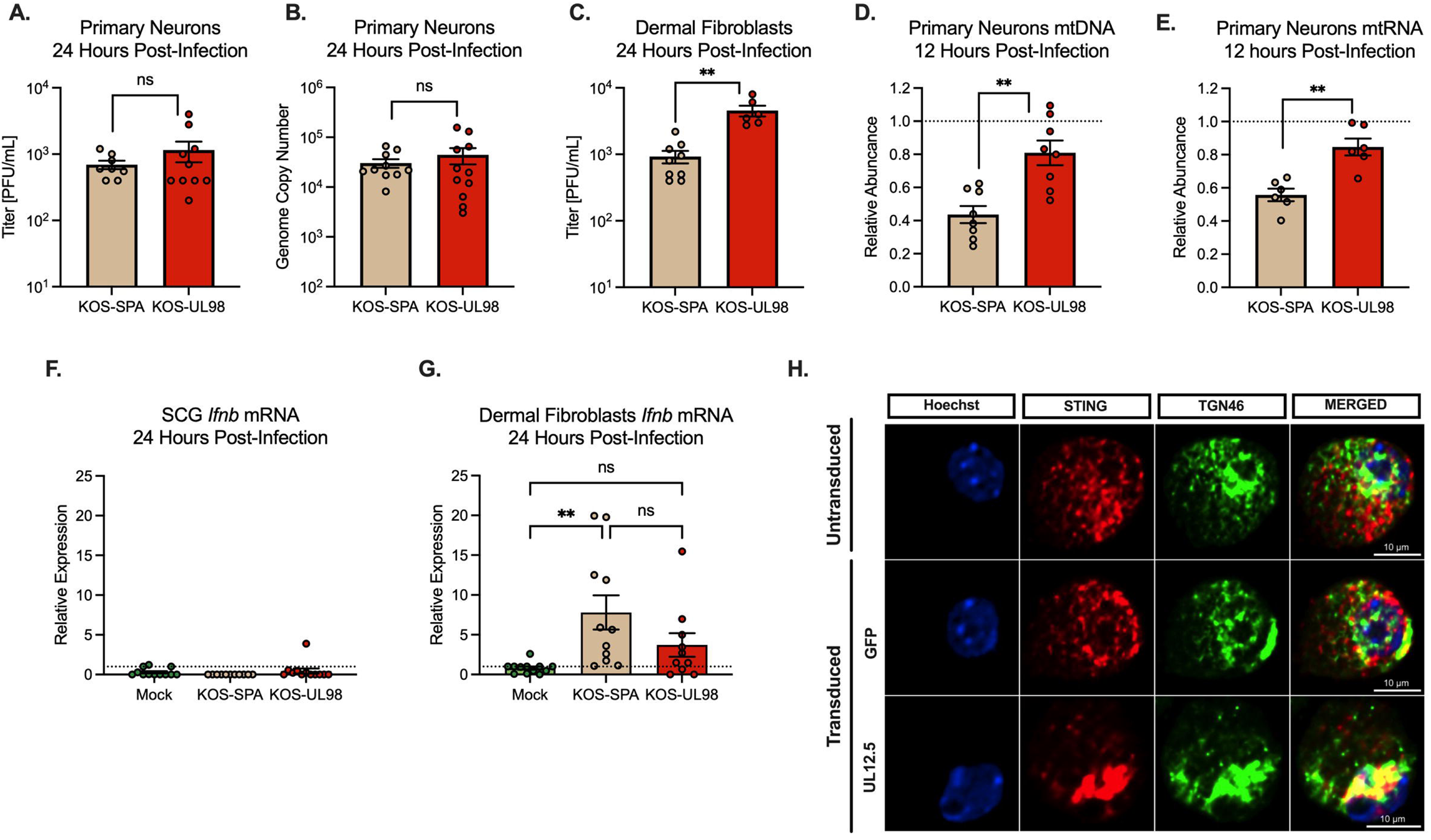
UL12.5 does not affect lytic replication in peripheral neurons. (A &B) Primary neurons isolated from the superior cervical ganglia (SCG) of newborn mice were infected at an MOI of 10 PFU/cell. Viral titer (A) and viral genome copy number quantified by qPCR (B) were measured at 24 hours post-infection with either KOS-SPA (WT) or KOS-UL98 (UL12.5 null). (C) Dermal fibroblasts isolated from newborn mice were infected at an MOI of 3 PFU/ml and viral titer quantified at 24 hours post-infection. (D & E) Relative abundance of (D) *mtDLoop1* mtDNA and (E) *mtCOX2* mtRNA transcripts measured by (RT-) qPCR 12 hours after infection with KOS-SPA or KOS-UL98. (F & G) Relative *Ifnb* mRNA expression 24 hours after infection of (F) primary sympathetic neurons or (G) dermal fibroblasts (H) Representative immunofluorescence images of STING/pIRF3 at 3-days post-transduction with either GFP or UL12.5 expressing lentiviral vector. All in A-G data are plotted as biological replicates from at least three independent experiments, shown as mean +/− SEM. Statistical comparisons were made using a t-test or one-way ANOVA with Tukey’s multiple comparison test. *p<0.05, **p<0.01

The absence of a discernible phenotype for expression of UL12.5 in HSV-1 replication following neuronal infection raised the possibility that UL12.5 functions differently in neurons compared to non-neuronal cells. One outcome of UL12.5 expression is decreased mitochondrial DNA and RNA ^27^, likely secondary to its release into the cytosol. Therefore, we carried out qPCR and RT-qPCR to evaluate levels of mitochondrial nucleic acids following infection with KOS-SPA or KOS-UL98. Importantly, we found that by 12 hours post-infection, both mtDNA and mtRNA were significantly decreased in peripheral neurons when infected with KOS-SPA compared to KOS-UL98 (Figure 1D and E). We also examined the downstream expression of interferon, focusing on *Ifnb.* In contrast to DF, HSV-1 infection of neurons did not result in a detectable increase in *Ifnb* expression (Figure 1F and G). We then validated that UL12.5 expression did result in the activation of nucleic acid sensing pathways in neurons by transducing neurons with a UL12.5-expressing lentivirus. For this experiment, UL12.5 was delivered using lentivirus-mediated transduction to specifically examine the effects of UL12.5. Neurons transduced with a UL12.5-expressing lentivirus displayed a marked increase in STING translocation to the Golgi compared to neurons transduced with a control GFP-expressing vector (Figure 1H), suggesting that UL12.5 expression alone is sufficient to activate STING in peripheral neurons. Therefore, UL12.5 is expressed during lytic infection of neurons and can activate STING pathways, but this does not result in the induction of *Ifnb* transcription, and neither restricts nor promotes HSV lytic replication.

### UL12.5 supports initial de-repression of lytic HSV genes following *in vivo* infection

Although UL12.5 had no significant role in productive replication in peripheral neurons, we hypothesized that UL12.5 might instead be supporting another phase of the HSV-1 life cycle specific to neurons, namely the establishment of or reactivation from a latent infection. To investigate the role of UL12.5 in latency and reactivation, we used a well-established mouse model of HSV-1 latent infection. Our goal was to investigate the ability of the UL12.5 mutant virus to reactivate from a latent infection. We infected mice with either KOS-SPA or KOS-UL98 via ocular scarification, which results in an acute infection in the peripheral ganglia, followed by the establishment of a latent infection ^52^. After latency had been established 28 days post-infection, the trigeminal ganglia (TG) were collected from infected mice, and viral genome copy numbers per ganglia were determined by qPCR. Interestingly, we found that KOS-UL98-infected TG had a significantly smaller pool of viral genomes at latency compared to those infected with SPA (Figure 2A). Therefore, UL12.5 expression supports the establishment of latency *in vivo* by promoting a larger pool of latent viral genomes.

**Figure 2.**
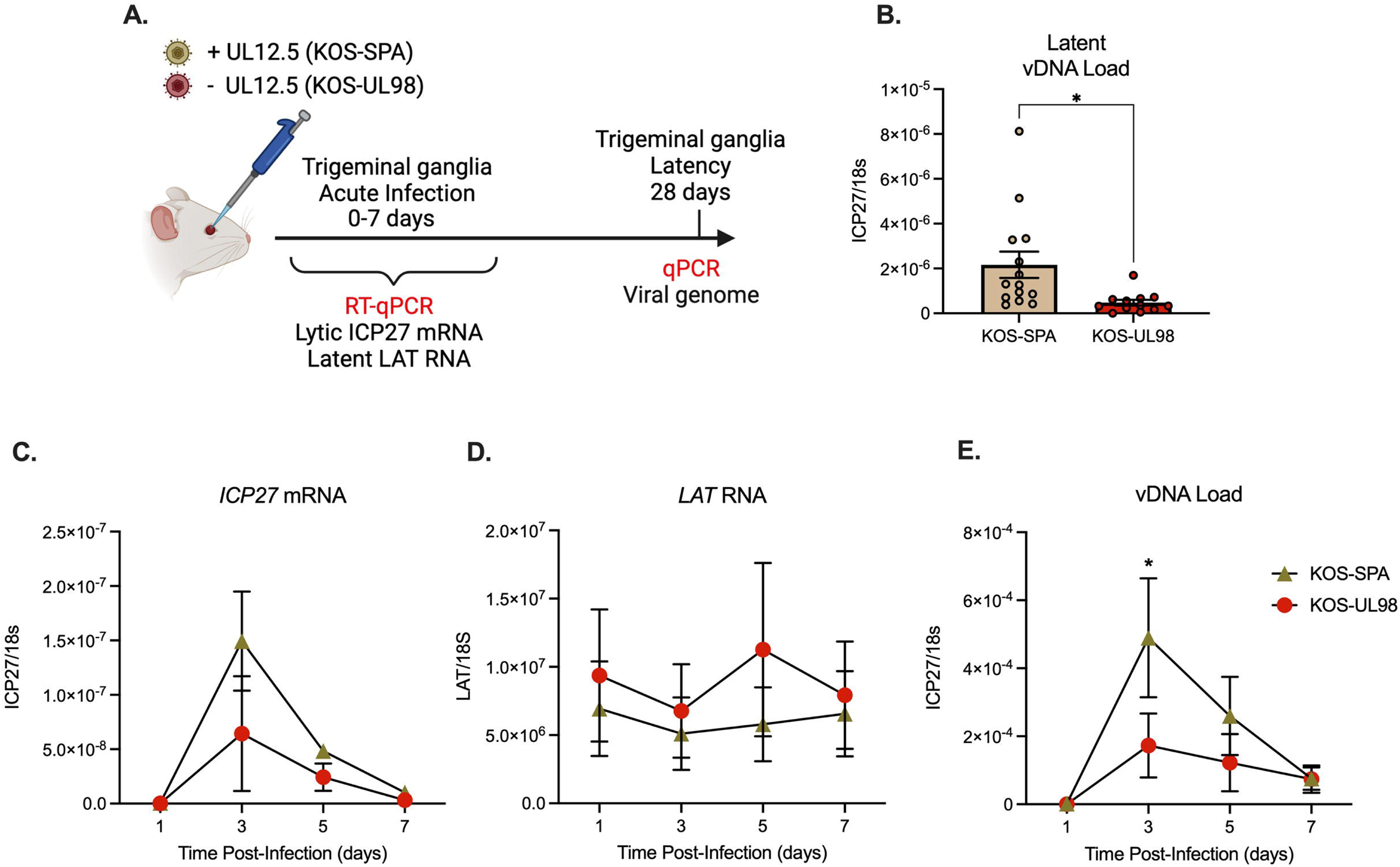
UL12.5 supports initial de-repression of lytic HSV genes following *in vivo* infection. (A) Schematic of the in vivo model of HSV-1 latent infection. (B) Quantification of the latent viral genome copy number measured by qPCR at 28 days after infection with KOS-SPA or KOS-UL98. The copy number of viral DNA was normalized to host 18s rDNA. (C & D) *ICP27* (C) or *LAT* (D) RNA copy numbers normalized to host 18S rRNA quantified by RT-qPCR during the acute infection period. (E) Viral genome copy number measured by qPCR over 7 days after infection with KOS-SPA or KOS-UL98 during the acute infection period. All data in B-E are plotted as biological replicates from at least three independent mouse experiments; shown as mean +/− SEM. Individual biological replicates are plotted in B. Statistical comparisons were made using a t-test (B) or one-way ANOVA with Tukey’s multiple comparison test (C-D). All statistical comparisons for time points in (C-E) are n.s. between viral strains except where otherwise noted. *p<0.05

The ability of HSV-1 to reactivate from latent infection *in vivo* is directly related to the latent viral DNA load ^53,54^. Therefore, we were unable to probe reactivation using this model. However, a previous study has shown that upon initial infection of the ganglia, HSV-1 undergoes a repression of lytic gene expression, which is then overcome, resulting in active lytic gene expression in some infected neurons to promote entry into the acute stage of infection ^55^. Therefore, we hypothesized that UL12.5 expands the pool of latent genomes by promoting the de-repression of viral genomes early during neuronal infection. To investigate the effect of UL12.5 on lytic gene expression during the establishment of latency *in vivo*, we infected mice with either KOS-SPA or KOS-UL98 via ocular scarification and quantified lytic and latent RNA transcripts, along with viral genome copy number, in TG early after infection. The expression of a representative viral lytic transcript (ICP27) was decreased in TG at early time points in mice infected with KOS-UL98 compared to those infected with KOS-SPA, although this decrease was not significant (Figure 2C). We did observe a statistically significant decrease in the viral DNA copy number at the peak of DNA replication (3 days post-infection) following infection with KOS-UL98 compared to KOS-SPA (Figure 3E). In contrast, expression of the *Latency-Associated Transcript* (*LAT*), a long noncoding RNA highly expressed during latency, was increased in mice infected with KOS-UL98 (Figure 2D). Thus, our data support the role of UL12.5 in promoting the initial de-repression of lytic genes early during infection *in vivo*.

**Figure 3.**
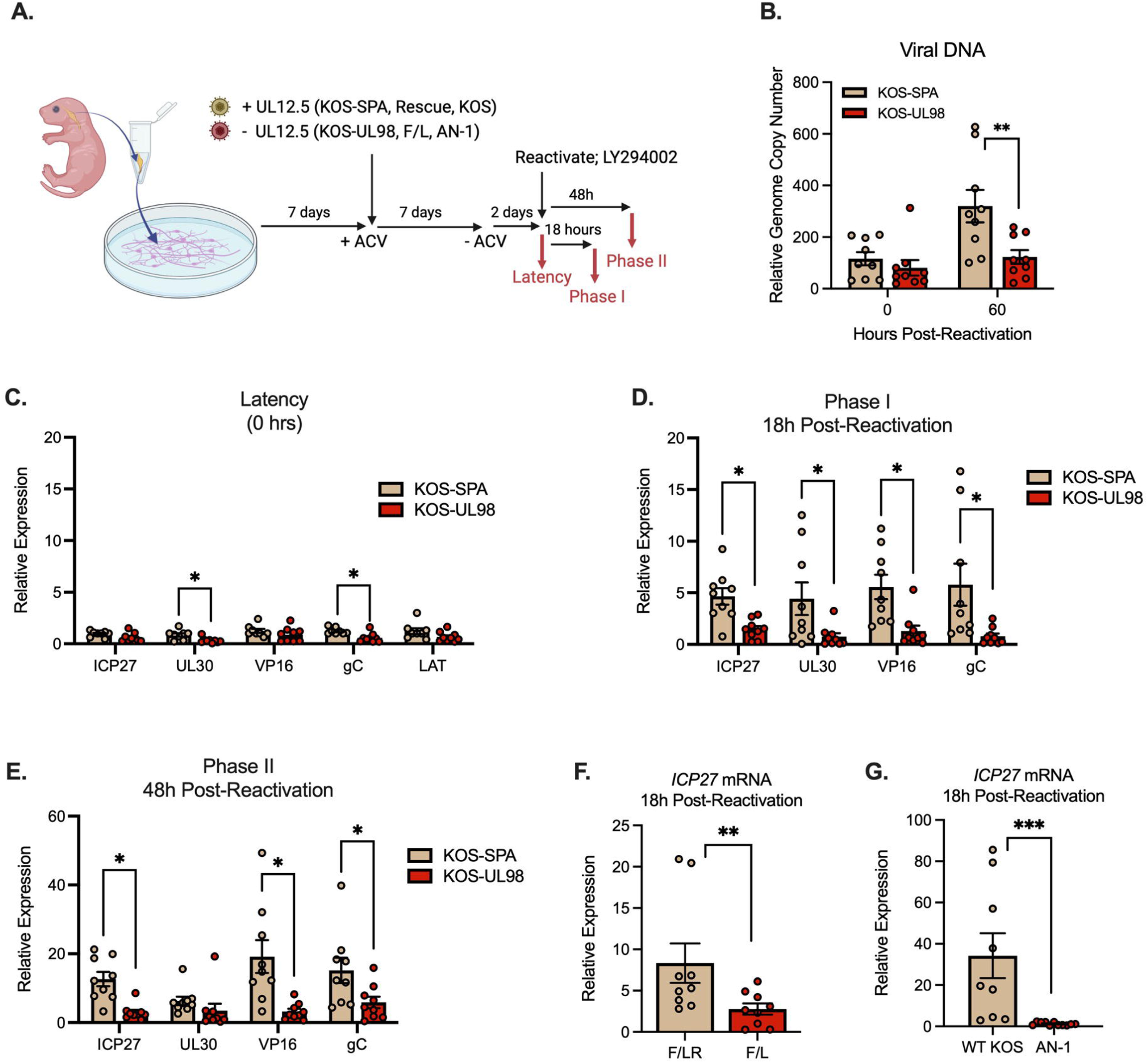
UL12.5 is required for Phase I lytic gene expression during HSV-1 reactivation. (A) Schematic of the in vitro HSV latent infection and reactivation model. Neuronal infection was carried out in the presence of acyclovir (ACV; 50 μM). Reactivation was induced using LY294002 (20 μM) in the presence of WAY-150138 (20 μM) to limit cell-to-cell spread. (B) Quantification of relative viral genome in latently infected (0 hours) and reactivated (60 hours) neurons infected with either KOS-SPA and KOS-UL98 measured by qPCR. (C) Quantification of lytic mRNAs and *LAT* in neurons latently infected with either KOS-SPA and KOS-UL98 measured by RT-qPCR for *ICP27* (IE), *UL30* (E), *VP16* (L), and *gC* (L). (D) Quantification of lytic mRNAs at 18 hours post-reactivation (Phase I). (E) Quantification of lytic mRNAs at 48 hours post-reactivation (Phase II). (F) Quantification of ICP27 mRNA during Phase I reactivation in neurons latently infected with F/L or the recue virus; F/LR. (G) Quantification of ICP27 mRNA during Phase I reactivation in neurons latently infected with AN-1 or the wild-type strain KOS. All in B-G data are plotted as biological replicates from at least three independent experiments; shown as mean +/− SEM. Statistical comparisons were made using a t-test (F-G) or one-way ANOVA with Tukey’s multiple comparison test. *p<0.05, **p<0.01, ***p<0.001

### UL12.5 is required for optimal lytic gene expression during HSV-1 reactivation

Based on our data suggesting that UL12.5 may support the derepression of the viral genome in neurons, we hypothesized that it may be important during the reactivation of HSV-1 from a latent infection. However, we were unable to test this *in vivo* because any effects on reactivation would be masked by the decreased latent genome copy number for the UL12.5 null virus. Instead, we turned to our *in vitro* model of HSV latency and reactivation (Figure 3A) ^32,34^. We isolated primary neurons from the SCG of postnatal mice and infected them with either KOS-UL98 or KOS-SPA in the presence of acyclovir (ACV). ACV functions to prevent viral DNA replication, and we have shown previously that ACV promotes latency establishment but does not impact reactivation ^36^. The benefit of using ACV in this experiment is that latent infection can be established with equivalent copy numbers of KOS-UL98 and KOS-SPA genomes. We confirmed that latency was established with equivalent DNA loads for KOS-UL98 and KOS-SPA (Figure 3B). Reactivation was triggered by the addition of a PI3 kinase (PI3K) inhibitor (LY294002; 20 μM), which is a well-established trigger of HSV-1 reactivation that mimics the loss of nerve growth factor signaling ^49^. Despite an equivalent amount of latent viral DNA, viral DNA replication following reactivation was decreased in KOS-UL98-infected neurons by 60 hours post-trigger compared to those infected with KOS-SPA (Figure 3B). We also compared the changes in viral DNA load between 0h and 60h for each virus; we observed an average 2.7-fold and 1.5-fold increase in DNA copy number for KOS-SPA and KOS-UL98, respectively. These data indicate that KOS-UL98 is unable to efficiently reactivate from a latent infection.

Viral DNA replication at 48h post-reactivation is a measurement of the later Phase II of reactivation. To determine whether UL12.5 is required for maximal Phase I or Phase II lytic gene expression, we quantified representative viral lytic mRNAs *ICP27*, *UL30*, *VP16* and *gC* at 0h (latent)18h (Phase I) and 48h (Phase II of reactivation). We observed a slight decrease in lytic gene expression during latent infection for *UL30* and *gC*, but not *ICP27* or *VP16* in the KOS-UL98 infected neurons; this may reflect low levels of spontaneous leaky viral gene expression (Figure 3C). We also quantified *LAT* expression as an indication of latency establishment following infection with the two different viruses and observed no change in LAT expression. Importantly, we found that the expression of all lytic genes tested was diminished during Phase I in the KOS-UL98 infected cultures (Figure 3D). We also observed significantly diminished expression of *ICP27*, *VP16*, and gC during Phase II of reactivation (Figure 3E). At this time point, *UL30* was less robustly expressed for KOS-SPA, and therefore, we did not detect a significant difference between the two viruses. These data suggest that UL12.5 plays a role in promoting Phase I gene expression.

To confirm the requirement of UL12.5 for lytic gene expression during Phase I of reactivation, we assessed reactivation from peripheral neurons using two additional HSV UL12.5 mutant viruses. The KOS F/L mutant contains mutations in the two potential translational start sites for UL12.5 that are associated with reduced expression of UL12.5 during lytic infection ^29^. This mutant has been used less frequently than KOS-UL98 because it still retains some capability to degrade mtDNA, albeit less robustly and at later time points than the wild-type ^29^. This is likely via an alternative mechanism to produce UL12.5 during lytic replication, such as truncation from full-length UL12. AN-1 is a UL12/UL12.5 deletion virus ^56^. Neurons infected with either KOS-F/L or AN-1 had decreased lytic gene expression during Phase I compared to their respective controls, the F/L recue virus (FLR) and wild-type (WT) KOS, when measured by RT-qPCR (Figure 3G & G). These results indicate that UL12.5 is required for Phase I gene expression during HSV-1 reactivation.

### UL12.5 is expressed and acts during Phase I to promote lytic gene expression

A previous study suggested that Phase I gene expression was independent of viral protein synthesis ^33^. This was based on the addition of cycloheximide (CHX), a protein synthesis inhibitor, to reactivating neurons *in vitro* at a time point prior to detectable lytic mRNA expression (10 hours post-trigger). Our data support a role for UL12.5 during Phase I, suggesting that the protein itself is synthesized following a reactivation trigger to promote later viral gene expression. However, an alternative was that UL12.5 is important during latency establishment to mediate later Phase I reactivation. To directly probe the function of UL12.5 during Phase I, we readdressed the impact of cycloheximide treatment on Phase I gene expression. Primary neurons were latently infected with a virus that is essentially wild-type but defective in cell-to-cell spread and expressing functional Us11-GFP to monitor full reactivation (Stayput-GFP) ^36,57^. Reactivation was induced, and cycloheximide was added either at the time of reactivation (0h) or 10 hours post-reactivation to inhibit protein synthesis. Using RT-qPCR to measure viral transcripts, we found that inhibition of protein synthesis at the start of reactivation (0hrs) restricted Phase I lytic gene expression (Figure 4A & B). We also confirmed that protein synthesis is not required after 10 hours post-trigger for maximal Phase I gene expression. These data suggest that viral and/or host protein synthesis within the first 10 hours after a reactivation trigger contributes to viral gene expression during Phase I, making it possible that UL12.5 itself may need to be translated for Phase I gene expression.

**Figure 4.**
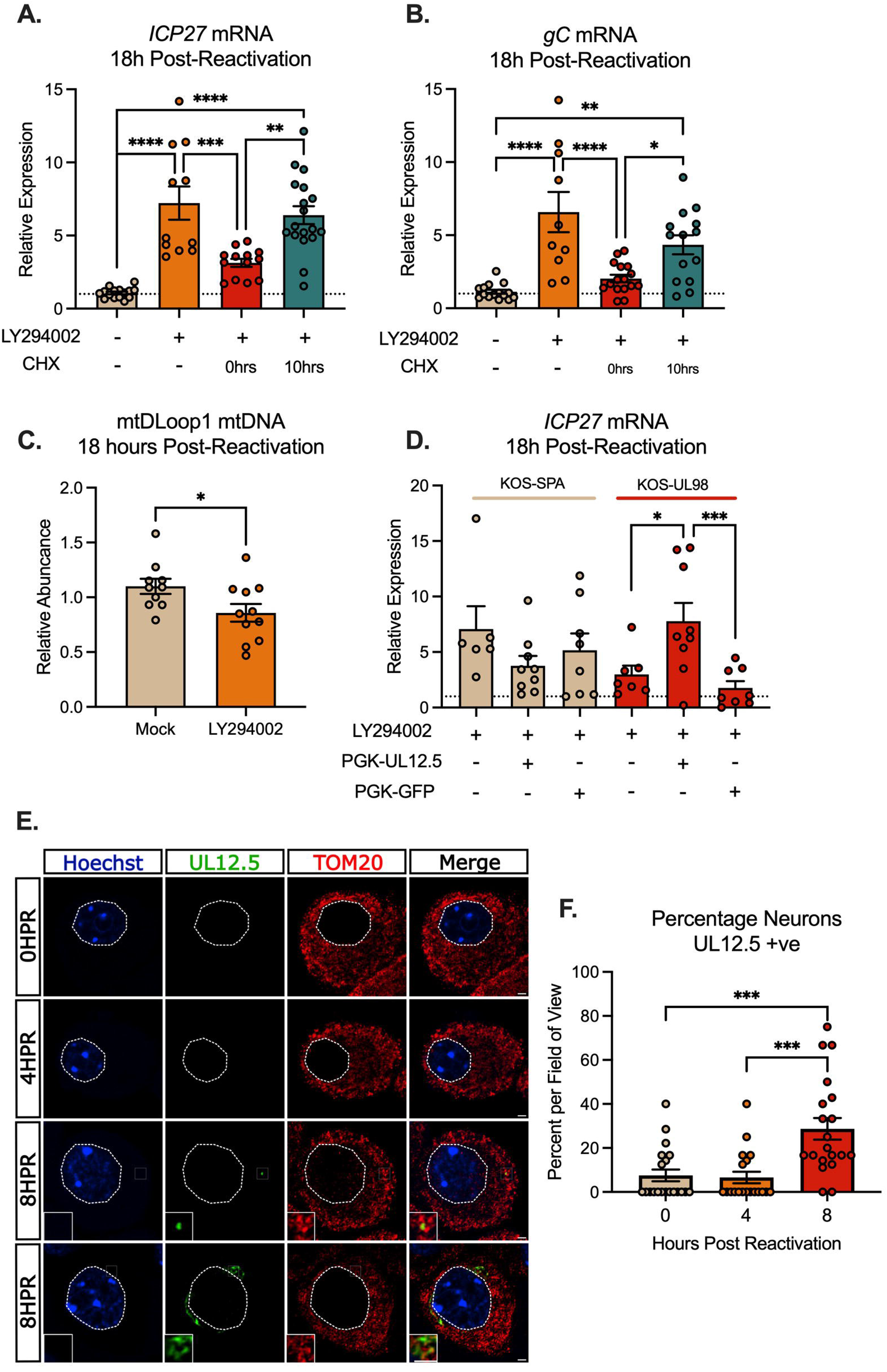
UL12.5 is expressed and functions during Phase I of HSV-1 reactivation. (A-B) Neurons were latently infected with HSV-1 Stayput-GFP and reactivated in the presence of cycloheximide (CHX; 10 μg/ml) added at either the same time as LY294002 or 10 hours later. Relative *ICP27* (A) and *gC* (B) mRNA was quantified by RT-qPCR. (C) Quantification of mtDNA abundance by qPCR of mtDLoop1 at 18 hours after reactivation post-reactivation. (D) Neurons latently infected with either KOS-SPA or KOS-UL98 were transduced with lentivirus expressing either GFP or UL12.5. The relative expression of *ICP27* mRNA was measured by RT-qPCR at 18 hours post-reactivation. (E) Representative immunofluorescence images of UL12/UL12.5 and TOM20 following reactivation. (F) Quantification of UL12.5 expressing neurons as a percentage of positive neurons per field of view at 0-, 4-, and 8 hours after reactivation. All in A-F data are plotted as biological replicates from at least three independent experiments; shown as mean +/− SEM. Statistical comparisons were made using a t-test (C) or one-way ANOVA with Tukey’s multiple comparison test. *p<0.05, **p<0.01, ***p<0.001, ****p<0.0001

Since UL12.5 depletes mtDNA and mtRNA during lytic infection of peripheral neurons, we hypothesized that if UL12.5 is expressed during Phase I, we may see a decrease in mtDNA. Therefore, we again established a latent infection with Stayput-GFP and induced reactivation with PI3-kinase inhibition. We found that mtDNA copy number was significantly reduced at 18h post-reactivation compared to latency (Figure 4C). It is important to note that this assay represents a bulk analysis of neurons, of which only a few may express UL12.5. Therefore, the amount of mtDNA depletion by the end of Phase I in reactivating neurons may be more pronounced than what was detected in bulk cultures.

To confirm that UL12.5 was expressed in Phase I, we sought to detect UL12.5 expression using immunofluorescence (IF) throughout the early stages of Phase I; a timepoint at which no viral proteins have previously been detected following in vitro reactivation. Following the reactivation of latently infected cultures, we performed immunofluorescence using an antibody that recognized both UL12 and UL12.5 ^24,26,58^. UL12 and UL12.5 can be differentiated based on their subcellular localization since UL12 is primarily nuclear, whereas UL12.5 is mitochondrial. Notably, we observed staining that co-localized with mitochondria (based on TOM20 staining) at 8 hours post-reactivation (Figure 4E). The staining pattern varied from small puncta of UL12/UL12.5 signal to more pronounced staining, with a range of between 0 and 80% of neurons per field of view positive for UL12.5 staining at this time-point (Figure 4F). These data suggest that UL12.5 protein is detectable in reactivating neurons prior to robust lytic gene expression at 18 hours post-reactivation, the Phase I time-point.

To directly determine whether UL12.5 can promote Phase I gene expression, as opposed to functioning during the establishment of latency, we complemented KOS-UL98 latently infected neurons with exogenous UL12.5. This approach also has the benefit of determining whether the defect in the ability of the KOS-UL98 virus to enter Phase I is due to a difference between HSV-1 UL12 and HCMV UL98 unrelated to UL12.5 (although the experiments with KOS F/L in Figure 3 support a UL12.5 specific function). Neurons were latently infected with either KOS-SPA or KOS-UL98 and then transduced with either a UL12.5 or GFP-expressing lentivirus vector. Reactivation was induced three days post-transduction, and Phase I gene expression was quantified. Transduction of KOS-SPA-infected neurons resulted in a slight but nonsignificant decrease in Phase I gene expression (Figure 4D). This may result from a decreased copy number of mtDNA at the time of reactivation, suboptimal expression, or timing of UL12.5 expression. Importantly, transduction with the UL12.5, but not control GFP, expressing lentivirus was able to increase reactivation for the KOS-UL98 infected cultures. Therefore, UL12.5 is both expressed during Phase I reactivation and can directly promote Phase I gene expression, strongly supporting a direct role for UL12.5 in the earliest wave of gene expression following exit from latency.

### Innate immune activation promotes lytic gene expression during Phase I

It is well-established that UL12.5 expression results in the activation of nucleic acid-sensing pathways ^6,23,28^. When considering the potential functional contribution of UL12.5 that supports lytic gene expression in Phase I, we hypothesized that activation of nucleic sensing pathways by UL12.5 supports lytic gene transcription during Phase I. To test this hypothesis, we investigated whether cGAS-STING activation could directly promote the reactivation of latently infected neurons. Agonists for both cGAS (ISD) and STING (DMXAA, Poly(dA-dT)/LyoVec, ADU-S100) ^59–61^ triggered reactivation based on the numbers of Us11-GFP positive neurons (Figure 5A-D). While the various cGAS-STING agonists induced varying degrees of reactivation, likely due to differences in activation mechanism, DMXAA induced comparable levels of reactivation to the well-established trigger LY294002. The addition of Poly(I:C)/LyoVec, which can activate RNA-sensing pathways, also triggered reactivation, but within a much smaller population of latently-infected neurons (Figure 5E). Therefore, activation of the nucleic acid-sensing pathways are sufficient to initiate viral reactivation.

**Figure 5.**
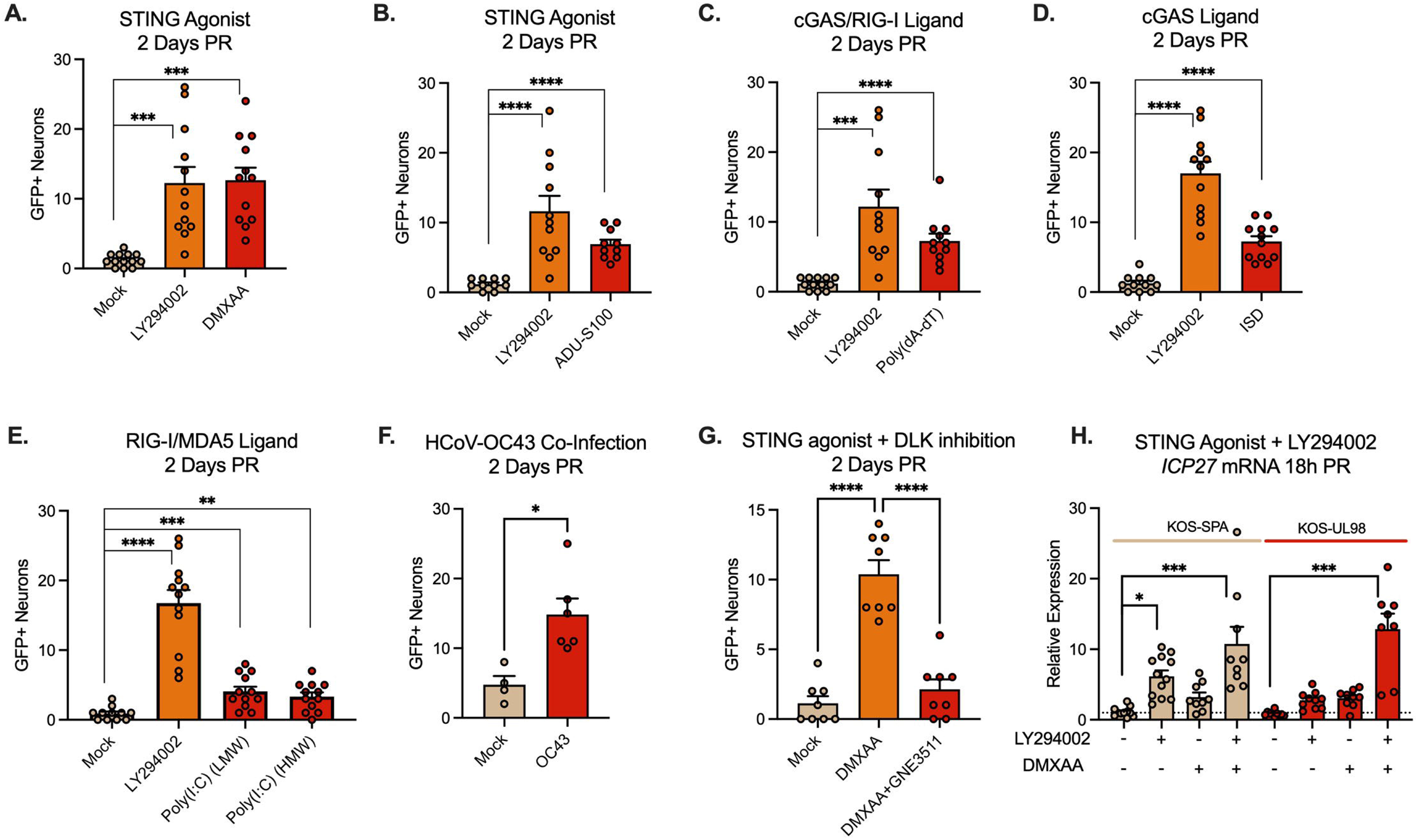
Activation of innate immune signaling induces HSV-1 reactivation from latent infection. (A-E) Neurons latently infected with Stayput-GFP were treated with ligands for the DNA/RNA sensing pathways using DMXAA (50 μg/mL; A), ADU-S100 (10 μg/mL; B), Poly(dA-dT)/LyoVec (5 μg/mL; C), ISD (5μg/mL; D) or Poly(I:C) (HMW), or Poly(I:C) (LMW) (10 μg/mL; E). (F) Neurons latently infected with Stayput-GFP were co-infected with the HCoV-OC43 at an MOI of 3 PFU/cell. Reactivation was quantified based on the numbers of Us11-GFP expressing neurons at 2 days post co-infection. (G) Neurons latently infected with Stayput-GFP were reactivated with DMXAA in the presence or absence of the DLK inhibitor GNE-3511 (4 μΜ). (G) Neurons latently infected with KOS-SPA and KOS-UL98 were treated with either LY294002 or DMXAA or LY294002, followed 2.5 hours later by DMXAA. All in A-F data are plotted as biological replicates from at least three independent experiments; shown as mean +/− SEM. Statistical comparisons were made using a t-test (F) or one-way ANOVA with Tukey’s multiple comparison test. *p<0.05, **p<0.01, ***p<0.001, ****p<0.0001

An important implication for the activation of innate sensing pathways as a trigger for HSV-1 reactivation is that co-infections may also induce reactivation. To test a virus that is capable of infecting murine neurons, we used the human coronavirus (HCoV)-OC43 ^62^. The addition of OC43 to latently infected neurons did not result in obvious changes associated with cell death. However, we did observe a low level of significantly increased reactivation (Figure 5F), indicating that co-infections may be a novel trigger for HSV-1 reactivation from neurons.

The activation of DLK and its downstream mediator JNK are critical steps in reactivation associated with numerous triggers in vitro and ex vivo ^32,34,36,37,50^. We therefore tested whether DMXAA-induced reactivation was DLK-dependent; we focused on DMXAA as this gave the most robust reactivation. Therefore, we treated neurons latently infected with Stayput-GFP with DMXAA, or DMXAA and the DLK inhibitor GNE-3511 (Figure 5E). Using GFP fluorescence as a readout for reactivation, GNE-3511 blocked reactivation induced by DMXAA. This suggests that HSV reactivation downstream of cGAS-STING activation is reliant on DLK.

Given that STING activation triggers HSV-1 reactivation and is the primary result of UL12.5 expression, we hypothesized that STING activation by DMXAA will rescue lytic gene expression during Phase I in the virus lacking UL12.5 expression. To test this hypothesis, we first established a latent infection in peripheral neurons *in vitro* with either KOS-SPA or KOS-UL98. After the establishment of latency, neurons were treated with LY294002 alone, DMXAA alone, or LY294002, followed by DMXAA 2.5 hours later (Figure 5H). Using RT-qPCR to measure viral transcripts, we found that when DMXAA was added 2.5 hours after LY294002 treatment, lytic gene expression recovered relative to levels in SPA-infected neurons. Intriguingly, the levels of viral lytic gene expression in both KOS-SPA and KOS-UL98 following reactivation induced by DMXAA were equivalent, indicating that UL12.5 is less important for reactivation induced by the STING agonist. Together, this suggests that UL12.5 induces lytic gene expression via activation of nucleic acid sensing pathways, including the cGAS/STING DNA sensing pathway.

### Loss of both the DNA and RNA sensing pathway restricts Phase I gene expression during HSV-1 reactivation

If UL12.5 activates nucleic acid sensing pathways to promote Phase I of HSV-1 reactivation, we hypothesized that down-regulation of these pathways would restrict lytic gene expression. To test this, we initially investigated the contribution of an intact cGAS/STING pathway on the induction of Phase I gene expression since UL12.5 has been hypothesized to result in the activation of this pathway, and our data support STING activation following UL12.5 expression in neurons. Therefore, we established latent infection in primary neurons by infection with Stayput-GFP. At 6 days post-infection, neurons were transduced two independent lentiviruses expressing shRNAs that target *Sting* mRNA or a non-targeting control. However, Phase I gene expression was equivalent in the STING depleted compared to the shRNA-transduced neurons (figure 6A). *Sting* knockdown was validated at the mRNA level (Figure 6B). Therefore, depletion of STING did not prevent entry into Phase I gene expression.

**Figure 6.**
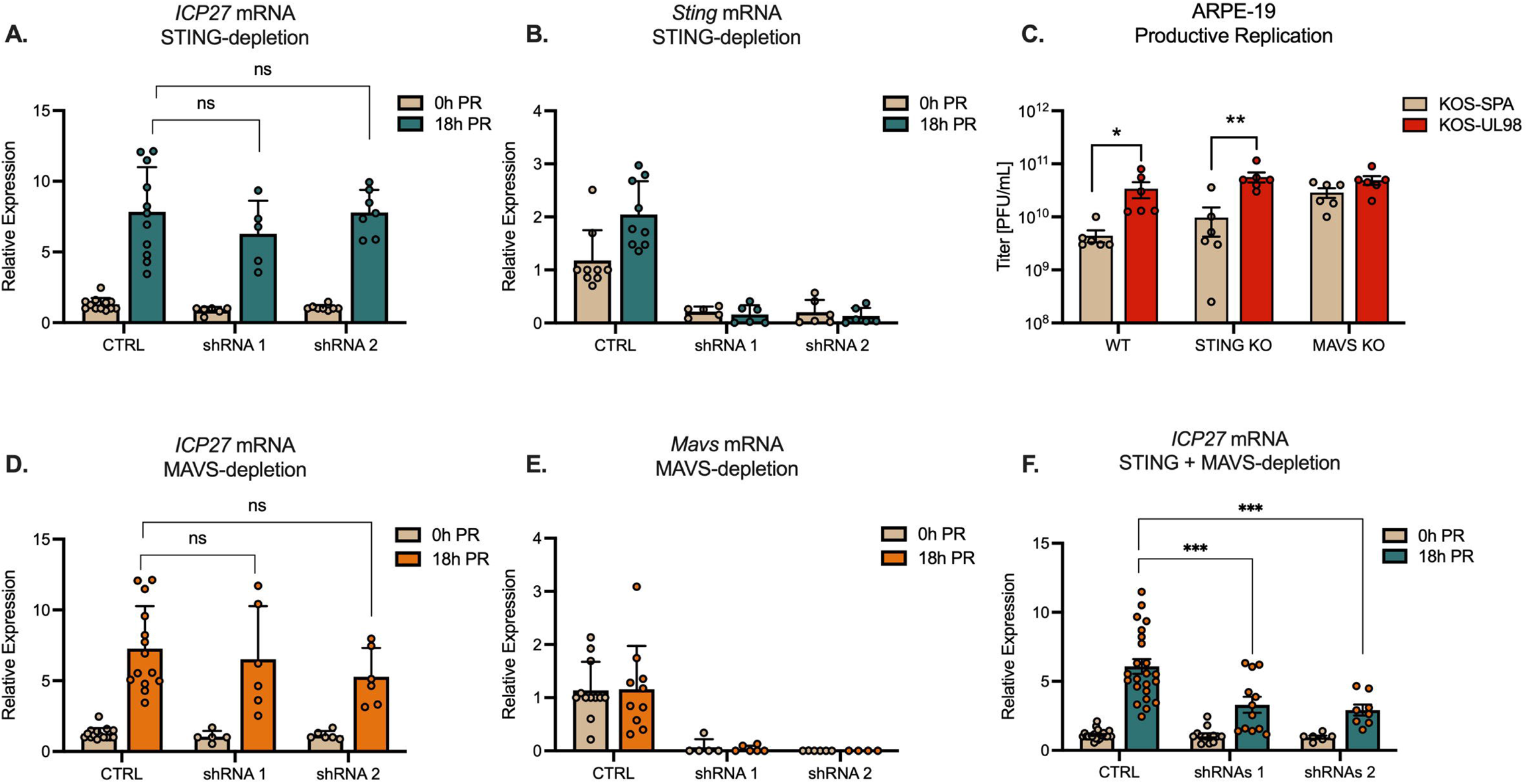
Intact STING and MAVS signaling is required for Phase I HSV-1 gene expression. (A & B) Latently infected neurons were transduced with vectors expressing shSTING or shCTRL during latency. Quantification of *ICP27 (A)* and *Sting* (B) transcripts by RT-qPCR at 0 and18 hours post-reactivation (PR) with LY294002 (20μM). (C) Titers of infectious virus at 24 hours post-infection of Wild-type, STING KO, or MAVS KO ARPE-19 cells infected with KOS-SPA or KOS-UL98 at an MOI of 5 PFU/cell. (D & E) Latently infected neurons were transduced with vectors expressing shMAVS or shCTRL during latency. Quantification of *ICP27 (D)* and *Mavs* (E) transcripts by RT-qPCR at 0 and 18 hours post-reactivation (PR) (F) Quantification of *ICP27* mRNA at 18 hours post-reactivation following depletion of both STING and MAVS. All in A-F data are plotted as biological replicates from at least three independent experiments; shown as mean +/− SEM. Statistical comparisons were made using a one-way ANOVA with Tukey’s multiple comparison test. *p<0.05, **p<0.01, ***p<0.001

UL12.5 expression can also result in the depletion of mtRNA, potentially as a result of release into the cytosol ^27^. Furthermore, a recent study suggested that UL12.5 expression can indirectly activate RNA sensing pathways via the activity of RNA polymerase III ^23^. Viruses that lack UL12.5 are known to replicate to higher levels in non-neuronal cells. However, the contribution of the RNA sensing pathway to this altered ability to replicate has not been investigated. Therefore, to determine the potential role of the MAVS-dependent RNA sensing pathway in the host response to UL12.5 expression, we first infected ARPE-19 cells with KOS-UL98 or KOS-SPA. ARPE-19s have previously been shown to lack DNA sensing pathways, including the cGAS/STING and ZBP1 sensing pathways. We found that KOS-UL98 still exhibited enhanced replication in ARPE-19s compared to KOS-SPA (Figure 6C). To confirm that this enhanced replication was via activation of the MAVS-dependent RNA sensing pathway, we infected previously described STING knock-out or MAVS knock-out ARPE-19 ^10^ cells with KOS-UL98 or KOS-SPA. Infection of the STING knock-out cells still resulted in enhanced replication of KOS-UL98 compared to KOS-SPA (Figure 6C), which was expected as this pathway is already deficient in these cells. However, replication of the two viruses was equivalent following infection of MAVS knock-out APRE-19s (Figure 6C). These data indicate that UL12.5 expression is capable of impacting HSV-1 viral replication via the MAVS-dependent RNA sensing pathway, and a lack of both DNA and RNA sensing pathways is required to prevent the UL12.5 mediated impact on replication in non-neuronal cells.

To determine whether the MAVS was required for HSV-1 reactivation, we then depleted MAVS from latently infected neurons and reactivated them as above. *Mavs* depletion was confirmed by RT-qPCR (Figure 6E). However, MAVS depletion did not impact Phase I gene expression (Figure 6D). Given the potential for UL12.5 to activate both DNA and RNA sensing pathways, we co-depleted both MAVS and STING from latently infected neurons. Depletion using two independent shRNA clones for each gene resulted in decreased Phase I lytic gene expression (Figure 6F). Therefore, inhibition of both the STING-dependent DNA sensing and MAVS-dependent RNA sensing pathways is required to prevent the exit of HSV-1 latency and entry into Phase I gene expression, suggesting that HSV can co-opt both the DNA and RNA sensing pathways to induce reactivation.

## Discussion

HSV-1 reactivation from a latent infection involves the activation of host cell pathways, with modification of AKT signaling, mTOR, and the DLK/JNK cell stress pathway, all linked to the virus’s reactivation ^33,49,63,64^. Here, we extend the known pathways to add activation of innate antiviral sensing pathways as a trigger for HSV-1 reactivation. Further, we show that even when reactivation is triggered by another mechanism (PI3-kinase inhibition), the virus uses its own protein, UL12.5, to activate antiviral sensing pathways and reactivate from a latent infection. This study has important implications for understanding the physiological triggers that can induce reactivation and identifying host proteins that mediate gene expression during reactivation. Further, the discovery that HSV-1 has a mechanism to activate host cell innate immune pathways during the earliest stage of reactivation is important to consider, given the life-long persistence of the virus in the nervous system.

Here, we found that UL12.5 is the first protein identified that is required for Phase I gene expression, which is the earliest wave of gene expression upon exit from latency ^33^. Phase I of HSV-1 reactivation is known to involve unique mechanisms of viral gene expression compared to productive replication in non-neuronal cells ^35^. In the initial discovery of Phase I by the Wilson lab, it was found that the key viral lytic transactivator, VP16, was not required for Phase I ^33^. VP16 is a tegument protein brought in with the virus during initial infection to promote the expression of immediate early genes. However, during the productive replication cycle, the *VP16* gene is expressed with late kinetics. During Phase I, *VP16*, like other late genes, is expressed in the absence of viral DNA replication ^33,36,37^. Further, work from the Sawtell and Thompson labs has shown that elements within the *VP16* promoter can function prior to immediate-early gene expression, specifically in neurons, and are involved in the initial depression of viral gene expression during acute infection of the ganglia ^55,65^. Viruses that lack VP16, in addition to the delivery of shRNA targeting of VP16 after latency has been established, have been used to demonstrate that VP16 is not required for Phase I gene expression in a primary neuronal model ^33^. An absence of a requirement for the main lytic transactivator for Phase I gene expression bolstered the hypothesis that Phase I gene expression is independent of viral protein and relies on the activation of host proteins. However, our data demonstrates a role for UL12.5 in promoting Phase I gene expression, showing that it acts even prior to the expression of a main lytic transactivator, VP16.

Primary neuronal systems of HSV-1 latency and reactivation have the advantage that pathways can be easily manipulated after latency has been established to uncover mechanisms of reactivation ^66^. The approach we used here, delivering UL12.5 into neurons to complement the reactivation defect of the KOS-UL98 virus, was important to validate a role for UL12.5, specifically during reactivation. Previous studies have found evidence of prior lytic gene expression in latently infected neurons ^67–69^. Further, our lab has found that altered conditions during latency establishment can impact the later ability of the virus to reactivate. For example, exposure to type I IFN during initial infection restricts later reactivation ^51^. Conversely, neurons exposed to increased cell stress during initial infection exhibit increased reactivation ^70^. Therefore, to fully determine that a viral protein functions specifically during reactivation, it is necessary to either perform a complementation experiment where a viral gene is reintroduced prior to reactivation or knock-out or known-down the viral gene after latent infection has been established.

Studies from our lab and others mapping the time of lytic gene expression during Phase I have found the peak of gene expression for key lytic transcripts like *ICP27*, *UL30*, and *VP16* to occur between 15- and 20 hours post-stimulus ^33,34,36^. The reason for the long delay in robust lytic gene expression following a reactivation stimulus is unknown. HSV-1 gene expression initiates from a silenced, heterochromatinized viral genome during reactivation ^71–76^; therefore, there are more obstacles to gene expression than *de novo* infection. However, this does still not perhaps explain the approximate 18-hour window from stimulus to lytic gene expression. Our data here now divides Phase I into two parts: the first 10-hour window requires protein synthesis, and the second part is independent of protein synthesis. Intriguingly, in a previous study investigating co-localization of the viral genome with a histone methyl/phospho switch during Phase I, we identified two waves of co-localization: one around 5 hours post-stimulus and one at 20 hours ^32^. Therefore, there could be an additional induction of lytic gene expression, at least for some viral transcripts, before 10 hours post-trigger. By bulk analysis of RNA, we were unable to detect robust UL12.5 mRNA (data not shown); however, this does not rule out the possibility of pulses of UL12.5 mRNA (and potentially other viral transcripts) in this early phase. In the future, higher-resolution single-cell techniques may be required to map viral transcription through Phase I.

So far, the host proteins known to function during Phase I to induce lytic gene expression are DLK and JNK ^32,34,36,37^. Experiments performed here with the DLK inhibitors suggest that DLK may be downstream of STING activation. However, an up-steam role in mediating UL12.5 expression cannot be ruled out, and it is possible that DLK functions at multiple levels during Phase I gene expression. In non-neuronal cells, activation of innate sensing pathways results in the activation of IRF3, NF-kB, and JNK. JNK activation results in the activation of the AP-1-containing protein, c-Jun. In neurons, however, JNK is constitutively active and has a physiological function in maintaining dendritic arborization ^77^. Following cell stress stimuli, JNK is redirected to induce c-Jun up-regulation and phosphorylation via DLK ^78,79^. It remains to be determined whether DLK can also function downstream of innate immune sensing pathways in neurons and whether UL12.5 potentially functions to maintain JNK activation for maximal Phase I gene expression. It also remains to be determined whether the activation of additional proteins, including IRF and NF-kB proteins, also play a role in HSV-1 reactivation. We have previously ruled out a role for c-Jun in mediating Phase I gene expression, although it is required to promote entry into full Phase II of reactivation^70^. JNK is not a transcription factor, nor does it have DNA binding capabilities. Therefore, additional factors likely function to stimulate lytic gene expression during Phase I. Our identification of a host innate sensing pathway in mediating Phase I gene expression will pave the way for multiple future studies to uncover proteins involved in HSV-1 reactivation and their mechanism of action.

Some of the best-defined reactivation triggers of HSV include psychological stress, fever, and sunburn ^31^. However, the majority of HSV reactivation events have no overt symptoms, and therefore, linking physiological triggers with reactivation can be problematic. Whether certain co-infections can reactivate latent HSV-1 *in vivo* remains to be determined, and analysis may be complicated based on the pro-or anti-viral effects of certain cytokines. Importantly, infection of neurons need not result in successful replication of the co-infecting agent, as our data suggest that the activation of innate sensing pathways, which would occur even following abortive infections, could promote reactivation. Further, the potential for endogenous, damage-associated molecules to induce HSV-1 reactivation is an important consideration. Mitochondrial dysfunction, particularly in the aging brain or neurodegenerating brain, can result in mitochondrial nucleic acid release and activation of downstream pathways sensing pathways ^80–82^. Enhanced reactivation of HSV-1 in this context may be an important consideration in disease pathogenesis.

A further implication of our study is the contribution of the RNA sensing pathway in detecting HSV infection, as we identified a role for MAVS in restricting HSV-1 replication in a UL12.5-dependent manner. MAVS is activated downstream of RNA sensing via RIG-I or MDA5. Ours is not the first to implicate sensing of host RNA in the response to HSV-1 infection. Work from the Gack lab has shown that the RNA sensor, RIG-I, can sense up-regulated host transcripts upon HSV-1 infection ^83^. Activation of RNA sensing pathways via cytoplasmic transcription by RNA polymerase III, including mtDNA, in macrophages has also been linked to innate immune responses to HSV-1 ^23,84^. The importance of RNA sensing pathways in HSV-1 infection is also highlighted by the ability of HSV-1 to limit immune activation mediated by the RNA sensing pathway, with most studies implicating HSV-1 in the inhibition of RIG-I activity ^85–88^. The potential mechanism of mtRNA release and degradation by UL12.5 is unknown. Likewise, the RNA sensor that may mediate the recognition of cytoplasmic mtRNA resulting from UL12.5 expression is unknown. There is some evidence that in macrophages, at least, RIG-I recognition could be via RNA polymerase III activity on mtDNA. However, the contribution of RNA polymerase III in non-macrophage cell types is unknown. Further, released mtRNA can directly activate RIG-I or MDA5; the exact sensor may vary depending on the nature of the released RNA ^10,11^. Therefore, it is possible that the direct release of mtRNA induces either RIG-I or MDA5 activation and subsequent MAVS activation to result in enhanced anti-viral responses in non-neuronal cells and promote Phase I reactivation in neurons.

STING signaling in neurons is still controversial and likely varies depending on the neuronal subtype and exposure of neurons to additional stimuli. For example, STING levels have been reported to be low in the majority of CNS neurons (with the exception being Purkinje neurons) ^89^. Our data would suggest that STING can be activated in sympathetic neurons but does not result in the upregulation of type I IFN. Direct STING activation in cultured in dorsal root ganglia cells has been shown to result in the release of type I IFN, which may either limit or enhance nociceptor firing capabilities and modulate the pain response ^90,91^. However, other studies have found limited type I IFN release with HSV-1 infection and in response to immune activation ^92–95^. Whether there are differences related to the nature of the stimulus, subtype of neuron, or the potential age of the mouse remains to be fully determined. It is important to note that the virus can establish latency in different subtypes of sensory and sympathetic neurons and can reactivate to varying degrees depending on the subtype ^96,97^. Further, certain stimuli, including cytokines, exposure to adjuvants, ischemia, and heat stress, can have been shown to increase the expression of components of nucleic acid sensing pathways, with the potential to enhance neuronal responses ^90,98,99^. The consequence of UL12.5 expression following exposure to different insults, in addition to in different neuronal subtypes, will be important to investigate to understand both the ability of the virus to reactivate from latency under different conditions and the impact on immune responses in the nervous system.

HSV-1 DNA during latent infection is nucleosomal and likely remains associated with nucleosomes at least through the earliest stages of reactivation ^71,100^. Because cGAS is tethered and inactivated by nucleosomes ^101–106^, the viral genome is hidden from the host foreign DNA detection mechanisms. Our data shows that UL12.5 can be expressed during initial exit from latency and result in mtDNA depletion, which implies that the reactivating neurons can now sense foreign DNA and RNA and mount a response. At the neuronal intrinsic level, the response does not appear to be anti-viral. Even if type I IFN is not released, there is evidence for cGAMP release and uptake into neighboring cells ^107^. There are also implications for both mitochondrial nucleic acid depletion and potential activation of innate immune responses, especially considering that there is evidence that Phase I gene expression can be abortive and not progress to full Phase II of reactivation ^33^. The potential long-term impact of UL12.5 expression in the context of full or abortive reactivation in different neuronal subtypes therefore warrants further investigation, especially given the potential links between HSV-1 infection and the development of late-onset Alzheimer’s disease.

### Limitations of the study

In our system, we were unable to show a direct role for mtDNA in HSV reactivation. Methods to deplete mtDNA include treatment with ethidium bromide or 2’,3’ dideoxycytidine. We attempted this in neurons, but at concentrations that were not toxic, the total copy numbers were still high even after long-term treatment. Another approach is to prevent mtDNA release. One mechanism of mtDNA release is via Bax/BAK pores. Sympathetic neurons primarily express Bax, and neuronal apoptosis is solely dependent on Bax ^108^. We have shown previously that HSV-1 reactivation is unaffected in Bax knock-out neurons ^34^. mtDNA can also be released via the mitochondrial permeability transition pore. We also found that reactivation was unaffected by blocking the port using cyclosporin. In the process of carrying out this study, Newmann et al. reported that mtDNA release following UL12.5 expression is independent of the Bax/BAK pore and the MPTP and instead uses an endosomal pathway involving Rab5 and Rab7 ^28^. However, inhibiting this pathway in latently infected neurons is problematic because the same proteins maintain NGF signaling via the TrkA signaling endosome^109^. A further minor weakness is that we used acyclovir to help promote latency establishment. However, we previously published that all the criteria for Phase I gene expression occur when latency is established in the absence of acyclovir ^36^. We did not use that system here because we wished to control for equivalent latent viral loads following infection with the UL12.5 mutant viruses.

## Materials and Methods

### Reagents

Compounds used in the study are described in Table 1. WAY-150138 was synthesized as described previously ^110^. Compound concentrations were used based on previously published IC50s and assessed for neuronal toxicity using cell body and axon health and degeneration index as previously published ^32^. All compounds used had an average score ≤1.

### Preparation of HSV-1 virus stocks

HSV-1 stocks of KOS-UL98, KOS-SPA, F/L, and F/L Rescue were propagated and titrated on Vero cells obtained from American Type Culture Collection (Manassas, VA). Vero cells were maintained in Dulbecco’s modified Eagle’s medium (Gibco) supplemented with 10% FetalPlex (Gemini Bio-Products). KOS-UL98, KOS-SPA, F/L, and F/L-Rescue viruses were kindly provided by James Smiley ^29^

Stocks of AN-1 and KOS37 (WT) were propagated and titrated on UL12 complementing 6-5 cells. 6-5 cells were maintained in Dulbecco’s modified Eagle’s medium (Gibco) supplemented with 10% FetalPlex (Gemini Bio-Products). KOS37 AN-1 and KOS37 (WT) and the complementing 6-5 cells ^56^ were kindly provided by Sandra Weller.

Stocks of Stayput Us11-GFP were propagated and titrated on gH-complementing F6 cells ^57^. Vero F6 cells were maintained in Dulbecco’s modified Eagle’s medium (Gibco) supplemented with 10% FetalPlex (Gemini Bio-Products) and 250μg/mL of G418/Geneticin (Gibco).

HCoV-OC43 stocks was obtained from ATCC (VR-1558) and was propagated and titrated on HCT-8 cells (ATCC CCL-244). Cells were maintained in Dulbecco’s modified Eagle’s medium (Gibco) supplemented with 10% Fetal Bovine Serum (Gibco) containing 5 unit/mL penicillin and 5 ug/mL streptomycin (Gibco).

### Mouse infections

Five-week-old female CD-1 mice (Charles River Laboratories) were anesthetized by intraperitoneal injection of ketamine hydrochloride (80 mg/kg of body weight) and xylazine hydrochloride (10 mg/kg) and inoculated with 1 × 10^6^ PFU/eye of virus (in a 5-μL volume) onto scarified corneas, as described previously ^74^. Mice were housed in accordance with institutional and National Institutes of Health guidelines on the care and use of animals in research, and all procedures were approved by the Institutional Animal Care and Use Committee of the University of Virginia. Mice were randomly assigned to groups, and all experiments included biological replicates from independent litters.

### Isolation of primary murine cells

Sympathetic neurons from the Superior Cervical Ganglia (SCG) of post-natal day 0-2 (P0-P2) CD1 Mice (Charles River Laboratories) Rodent handling and husbandry were carried out under animal protocols approved by the Animal Care and Use Committee of the University of Virginia (UVA). Ganglia were briefly kept in Leibovitz’s L-15 media with 2.05 mM l-glutamine before dissociation in collagenase type IV (1 mg/ml) followed by trypsin (2.5 mg/ml) for 20 min; each dissociation step was at 37°C. Dissociated ganglia were triturated, and approximately 10,000 neurons per well were plated onto rat tail collagen in a 24-well plate. Sympathetic neurons were maintained in feeding media: Neurobasal® Medium supplemented with PRIME-XV IS21 Neuronal Supplement (Irvine Scientific) or MACS® NeuroBrew®-21 (Miltenyi Biotec), 50 ng/ml Mouse NGF 2.5S, 2 mM l-Glutamine, and Primocin). Aphidicolin (3.3 µg/ml) was added to the media for the first five days post-dissection to select against proliferating cells.

For isolation of primary murine dermal fibroblasts skin was cut from the back of the neck of decapitated mouse pups and digested with collagenase type IV (1 mg/mL) and trypsin (2.5 mg/mL) with rotation (50 rpm) for 1 hour at 37°C. To the mixture was triturated and the cell suspension was filtered through Corning® 100µm cell strainer and cells cultured in Dulbecco’s modified Eagle’s medium (DMEM; Gibco) supplemented with 10% fetal bovine serum (FBS; Gibco) and Primocin (InvivoGen; 100 µg/ml).

### Lytic HSV-1 infections

Post-natal day 6-8 neurons were infected with UL98/SPA at MOI 10 PFU/cell, (assuming 10,000 cells per well) in Dulbecco’s Phosphate Buffered Saline (DPBS) + CaCl2 + MgCl2 supplemented with 1% fetal bovine serum and 4.5 g/L glucose for 3.5 h at 37°C. Primary murine Dermal Fibroblast were infected with UL98/SPA at MOI 3 PFU/cell, (assuming 50,000 cells per well) in inoculation medium (as described above) on a shaker (50 rpm) for 1h at 37°C. The inoculum was replaced with feeding media (as described above for neurons) or Dulbecco’s modified Eagle’s medium (Gibco) supplemented with 1% Fetal Bovine Serum (Gibco) for Dermal Fibroblast. The hTERT wilt-type and STING or MAVS knock-out ARPE-19 cells have been described previously^10^. Cells were infected with KOS-SPA or KOS-UL98 at MOI of 5 PFU/cell and inoculum was replaced with feeding media as described above for Dermal Fibroblasts. For quantification of infectious virus, both the cells and supernatant were collected, and samples were freeze-thawed three times to lyse cells. Homogenates were titrated on a monolayer of Vero cells.

### Establishment and reactivation of latent HSV-1 infection in primary neurons

Post-natal day 6-8 neurons were infected with Stayput US11-GFP at MOI 10 PFU/cell, KOS-UL98/KOS-SPA at MOI 10 PFU/cell, F/L/F/L-Rescue at MOI 5 PFU/cell, or AN-1/WT at MOI 5 PFU/cell (assuming 10,000 cells per well) in Dulbecco’s Phosphate Buffered Saline (DPBS) + CaCl_2_ + MgCl_2_ supplemented with 1% fetal bovine serum, 4.5 g/L glucose, and 10 mM acyclovir (ACV) for 4 h at 37°C. Post-infection, the inoculum was replaced with feeding media (as described above) with 50 mM ACV. 7 days post-infection, ACV was washed out and replaced with feeding media alone. Reactivation was quantified by counting the numbers of GFP-positive neurons or performing reverse transcription–quantitative PCR (RT–qPCR) of HSV-1 lytic mRNAs isolated from the cells in culture. WAY-150138 (10 µg/ml) was added following ACV wash-out. Reactivation induced by LY294002 (20 µM), or the immunostimulatory agents DMXAA (50 µg/mL), ADU-S100 (10 µg/mL), ISD (5 µg/mL), Poly(dA:dT)/LyoVec (5 µg/mL), High Molecular Weight Poly(I:C) (HMW) (10 µg/mL) or Low Molecular Weight Poly(I:C)(LMW) (10 µg/mL).

### Analysis of mRNA expression by reverse transcription–quantitative PCR (RT– qPCR)

Total RNA was extracted from approximately 1.0 × 10^4^ neurons using the Quick-RNA™ Miniprep Kit (Zymo Research) with an on-column DNase I digestion. mRNA was converted to cDNA using the Maxima First Strand cDNA Synthesis Kit for RT-qPCR (Fisher Scientific), using random hexamers for first-strand synthesis and equal amounts of RNA (20–30 ng/reaction). To assess viral DNA load, total DNA was extracted from approximately 1.0 × 10^4^ neurons using the Quick-DNA™ Miniprep Plus Kit (Zymo Research). qPCR was carried out using PowerUp™ SYBR™ Green Master Mix (ThermoFish Scientific). The relative mRNA or DNA copy number was determined using the comparative CT (ΔΔCT) method normalized to mRNA or DNA levels in latently infected samples. Primers used are shown in Table S2. Viral RNAs were normalized to mouse reference gene 18S rRNA. All samples were run in triplicate on an Applied Biosystems™ QuantStudio™ 6 Flex Real-Time PCR System and the mean fold change compared to the calculated reference gene. Exact copy numbers were determined by comparison to standard curves of known DNA copy number of viral genomes.

### Preparation of lentiviral vectors

Lentiviruses expressing shRNA against STING and MAVS (STING-1 TRCN0000346321, STING-2 TRCN0000346266, MAVS-1 TRCN0000124769, MAVS-2 TRCN0000124772), or a control lentivirus shRNA (pLKO.1 vector expressing a non-targeting shRNA control) were prepared by co-transfection using JetPRIME® with psPAX2 and pCMV-VSV-G ^111^ into the 293LTV cell line (Cell Biolabs). psPAX2 was a gift from Didier Trono (Addgene plasmid # 12260 ; http://n2t.net/addgene:12260 ; RRID:Addgene_12260) and pCMV-VSV-G was a gift from Bob Weinberg (Addgene plasmid # 8454 ; http://n2t.net/addgene:8454 ; RRID:Addgene_8454). The same approach was used to generate lentiviruses expressing UL12.5-SPA (PGK-UL2.5-SPA), or a control lentivirus, expressing GFP (pLenti PGK-GFP-Puro ^112^). pLenti PGK GFP Puro (w509-5) was a gift from Eric Campeau & Paul Kaufman (Addgene plasmid # 19070 ; http://n2t.net/addgene:19070 ; RRID:Addgene_19070). The PGK-UL12.5-SPA plasmid was generated by cloning UL12.5-SPA from pLenti CMV Neo UL12.5-SPA ^29^, which was a gift from Dr. James Smiley, into pCDH-PGK. pCDH-PGK was a gift from Kazuhiro Oka (Addgene plasmid # 72268 ; http://n2t.net/addgene:72268 ; RRID:Addgene_72268). Supernatant was harvested at 40- and 64-h post-transfection and filtered using a 45μM PES filter. Sympathetic neurons were transduced overnight in neuronal media containing 8μg/ml protamine sulfate and, for latently infected neurons, 50 μM ACV.

### Immunofluorescence

Neurons were fixed for 15 minutes in 4% Formaldehyde and blocked in 5% Bovine Serum Albumin and 0.3% Triton X-100 and incubated overnight in primary antibody. Antibodies and concentrations are described in Table S3. Following primary antibody treatment, neurons were incubated for one hour in Alexa Fluor® 488-, 555-, and 647-conjugated secondary antibodies for multi-color imaging (Invitrogen). Nuclei were stained with Hoechst 33258 (Life Technologies). Images were acquired using an sCMOS charge-coupled device camera (pco.edge) mounted on a Nikon Eclipse Ti Inverted Epifluorescent microscope using NIS-Elements software (Nikon). Images were analyzed and intensity quantified using ImageJ.

### Statistical analysis

Power analysis was used to determine the appropriate sample sizes for statistical analysis. All statistical analysis was performed using Prism V10. The normality of the data was determined with the Kolmogorov-Smirnov test. Specific analyses are included in the figure legends.

## Acknowledgments

This work was supported by the Owens Family Foundation (ARC), a UVA Global Infectious Disease Institute Seed award (ARC), UVA Wagner Fellowships (SAD and ALW), and National Institute of Health grants R21AI171544 (ARC), T32AI007046 (SRC and ALW), T32GM008136 (SAD), R01AG085782 to Dr. Agnel Sfeir, Memorial Sloan Kettering Cancer Center (to support MT).

We thank Dr. James Smiley (University of Alberta) for the KOS-UL-98, KOS-SPA, F/L, and F/LR viruses, along with the pLenti CMV Neo UL12.5-SPA plasmid. We thank Dr. Sandra Weller (University of Connecticut) for the AN-1 virus, the complementing 6-5 cells and the UL12/UL12.5 antibody. We thank Gary Cohen (University of Pennsylvania) for the Vero F6 cells.

## Author contributions

SRC: Investigation, writing – original draft, visualization, formal analysis.

MEF: Investigation, validation, writing – original draft, visualization, formal analysis.

PAK : Investigation, validation, writing – original draft, visualization.

ALW: Investigation, visualization, writing – reviewing & editing, formal analysis.

SAD: Investigation, resources.

AB: Investigation.

TM: Resources.

MT: Resources

DAE: Resources, writing – reviewing & editing.

ARC: Conceptualization, methodology, investigation, writing – original draft, writing – reviewing & editing, formal analysis, visualization, supervision, project administration, funding acquisition.

