## Supplemental table 1 and table 2 for "Co-option of mitochondrial nucleic acid sensing pathways by HSV-1 UL12.5 for reactivation from latent Infection"

Table S1: Compounds Used and Concentrations

| **Table 1. Compounds used and concentrations** | | | |
| --- | --- | --- | --- |
| **Compound** | **Supplier** | **Identifier** | **Concentration** |
| Aphidicolin (APH) | AG Scientific | A-1026 | 3.3 µg/ml |
| Acycloguanosine | Millipore Sigma | A4669 | 10 μM, 50 μM |
| Cycloheximide | Sigma-Aldrich | C4859 | 10 μg/mL |
| LY294002 | Tocris | 1130 | 20 µM |
| G418/Geneticin | Gibco | 10131-035 | 250μg/mL |
| WAY-150138 | Pfizer | N/A | 10 μg/mL |
| DMXAA | Invivogen | Tlrl-dmx | 50 μg/mL |
| ISD/LyoVec™ | Invivogen | tlrl-isdc | 5 μg/mL |
| ADU-S100 | Invivogen | tlrl-nacda2r-01 | 10 μg/mL |
| Poly(I:C) LMW/LyoVec™ | Invivogen | tlrl-picwlv | 10 μg/mL |
| Poly(I:C) HMW/LyoVec™ | Invivogen | tlrl-piclv | 10 μg/mL |
| Poly(dA:dT)/LyoVec™ | Invivogen | tlrl-patc | 5 μg/mL |
| GDNF | PeproTech | 450-44 | 50 ng/ml |
| NGF 2.5S | Alomone Labs | N-100 | 50 ng/mL |
| MACS® NeuroBrew®-21 | Miltenyi Biotec | 130-093-566 | 1x |
| PRIME-XV IS21 Neuronal Supplement | Irvine Scientific | 91142 | 1X |
| L-glutamic acid | Millipore Sigma | G5638 | 3.7 µg/ml |
| D-glucose | Sigma-Aldrich | G8769-100ML | 4.5 g/L |
| Fetal Bovine Serum (FBS) | Gibco | 26140079 | 10% or 1% |
| FetalPlex | Gemini Bio-Products | 50-753-2987 | 10% |
| Primocin | Invivogen | ant-pm-1 | 100 µg/ml |
| Penicillin-Streptomycin | Gibco | 15070063 | 5 unit/mL - 5 ug/mL |
| Trypsin | Worthington Biochemical | LS004454 | 1x |
| Dulbecco's phosphate-buffered saline (DPBS) | Gibco | 14190144 | 1X |
| DPBS, calcium, magnesium | Gibco | 14040133 | 1X |
| DMEM high glucose | Gibco | 11965092 | 1X |
| Leibovitz’s L-15 Medium | Gibco | 11415064 | 1X |
| Neurobasal® Medium | Gibco | 12348017 | 1X |
| Bovine Serum Albumin (BSA) | Gold Biotechnology. Inc. | A-420-25 | 5% |
| jetPRIME | VWR | 89129-920 | 1X |
| Protamine sulfate | Sigma-Aldrich | P3369-10G | 8ug/ml |
| Methyl cellulose | Fisher Scientific | M352-500 | 1% |
| Triton-X | Thermofisher | X100-100ML | 0.3% |
| Formaldehyde Solution | Sigma-Aldrich | F8775-4X25ML | 4% |
| Hoechst 33258 Stain | Sigma-Aldrich | 94403 | 0.01% |
| ProLong™ Diamond Antifade Mountant | Life Technologies | P36970 | 1X |
| Odessey Blocking Buffer | LI-COR | 927-40000 | 1X |
| Maxima First Strand cDNA Synthesis Kit | Thermo Scientific™ | K1641 | 1x |
| Maxima Reverse Transcriptase | Thermo Scientific™ | EP0741 | 1X |
| Quick-DNA/RNA™ MiniPrep Plus Kit | Zymo Research | D7003 | 1X |
| PowerUp™ SYBR™ Green Master Mix | Thermo Fisher Scientific | A25742lif | 1X |

Table S2: Primers Used for RT-qPCR

| Primer | Sequence 5’ to 3’ |
| --- | --- |
| m 18S rRNA F | CAC GGA CAG GAT TGA CAG ATT |
| m 18S rRNAR | GCC AGA GTC TCG TTC GTT ATC |
| m Ifnb F | CCC TAT GGA GAT GAC GGA GA |
| m Ifnb R | CCC AGT GCT GGA GAA ATT GT |
| m MT-COX2 F | ATA ATC CCA ACA AAC GAC CT |
| m MT-COX2 R | CTC GGT TAT CAA CTT CTA GCA |
| m Dloop F | CCAGTCTTGTAAACCGGAGA |
| m Dloop R | CTATCACCCTATTAACCACTC |
| m Sting F | CCC AAC ATT CGA TTC CGA GAT A |
| m Sting R | CTC GTA GAC GCT GTT GGA ATA A |
| m MAVS F | TTA CGC CTG GGA GTC CGC AT |
| m MAVS R | TGG GAT GGA CTG AGA TGG ACT GC |
| ICP27 F | GCA TCC TTC GTG TTT GTC ATT CTG |
| ICP27 R | GCA TCT TCT CTC CGA CCC CG |
| UL30 F | CGC GCT TGG CGG GTA TTA ACA T |
| UL30 R | TGG GTG TCC GGC AGA ATA AAG C |
| VP16 F | GGA CCG GAC GGA CCT TAT |
| VP16 R | GGT TGC TTA AAT GCG TGG TG |
| gC F | GAG TTT GTC TGG TTC GAG GAC |
| gC R | ACG GTA GAG ACT GTG GTG AA |
| LAT F | TGT GTG GTG CCC GTG TCT T |
| LAT R | CCA GCC AAT CCG TGT CGG |

Table S3: Antibodies Used for Immunofluorescence

| Antibody | Supplier | Identifier | Concentration | RRID |
| --- | --- | --- | --- | --- |
| Mouse STING | Millipore Sigma | MABF270 | 1:50 | AB_3106861 |
| Rabbit TGN46 | Novus Biologicals | NBP1-49643 | 1:400 | AB_10011762 |
| Rabbit UL12/12.5 | Gift from Dr. Weller | BWp12 | 1:5000 | AB_3106862 |
| Mouse TOM20 | Abnova | H00009804-M01 | 1:200 | AB_1507602 |
| F(ab’)2 Goat anti Rabbit IgG (H+L) Alexa Fluor® 647 | Invitrogen | A21237 | 1:1000 | AB_1500743 |
| F(ab’)2 Goat anti Mouse IgG (H+L) Alexa Fluor® 555 | Invitrogen | A21246 | 1:1000 | AB_2535814 |
